# The neurokinin-1 receptor is expressed with gastrin-releasing peptide receptor in spinal interneurons and modulates itch

**DOI:** 10.1101/2020.07.14.199471

**Authors:** Tayler D. Sheahan, Charles A. Warwick, Louis G. Fanien, Sarah E. Ross

## Abstract

The neurokinin-1 receptor (NK1R, encoded by *Tacr1*) is expressed in spinal dorsal horn neurons and has been suggested to mediate itch. However, previous studies relied heavily on neurotoxic ablation of NK1R spinal neurons, which limited further dissection of their function in spinal itch circuitry. Thus, we leveraged a newly developed *Tacr1*^*CreER*^ mouse line to characterize the role of NK1R spinal neurons in itch. We show that pharmacological activation of spinal NK1R and chemogenetic activation of *Tacr1*^*CreER*^ spinal neurons increases itch behavior, whereas pharmacological inhibition of spinal NK1R suppresses itch behavior. We use fluorescence *in situ* hybridization to characterize the endogenous expression of *Tacr1* throughout the superficial and deeper dorsal horn, as well as the lateral spinal nucleus.

Retrograde labeling studies from the parabrachial nucleus show that less than 20% of superficial *Tacr1*^*CreER*^ dorsal horn neurons are spinal projection neurons, and thus the majority of *Tacr1*^*CreER*^ are local interneurons. We then use a combination of *in situ* hybridization and *ex vivo* two-photon Ca^2+^ imaging of the spinal cord to establish that NK1R and the gastrin-releasing peptide receptor (GRPR) are coexpressed within a subpopulation of excitatory superficial dorsal horn neurons. These findings are the first to describe a role for NK1R interneurons in itch and extend our understanding of the complexities of spinal itch circuitry.

## Introduction

Acute itch, much like pain, serves as a protective warning signal to the body, indicating that there is an irritant on the skin that needs to be removed. Itchy stimuli are first detected by primary afferent fibers innervating the skin, which then relay inputs to neurons within the spinal cord dorsal horn including local interneurons, as well as ascending spinal projection neurons that target brainstem, midbrain, and thalamic structures. However, how these spinal neurons process and encode itch as well as other somatosensory stimuli is not fully understood.

A major unresolved question regarding the spinal circuitry of itch and pain is how to distinguish spinal projection neurons from interneurons. Proposed distinguishing criteria include morphology or genetic markers [21,64]. For instance, the neurokinin-1 receptor (NK1R), an excitatory G-protein coupled receptor that is expressed throughout the nervous system and activated by the neuropeptide substance P (SP), is often highlighted as a selective marker for spinal projection neurons [9,18,25,28,64,71]. Yet *Tacr1*, which encodes NK1R, appears to be expressed within multiple excitatory dorsal horn neuron populations recently identified by single cell RNA-sequencing [25,70]. Thus, whether NK1R is a true marker of spinal projection neurons requires further investigation.

NK1R has been studied extensively as a putative target for pain relief in preclinical rodent models [40,43,66,68], but demonstrated negligible clinical success and was quickly dropped as a therapeutic target [14,26]. Intriguingly, NK1R expressed on spinal neurons has recently emerged as a potential therapeutic target for chronic itch. Neurotoxic ablation studies show that loss of NK1R spinal neurons reduces scratching in rodents [1,3,19]. However, the effect of modulation of NK1R spinal neurons in an intact spinal circuit remains unclear. Furthermore, little is known about which NK1R neurons are involved in itch processing or where they fit within the current model of itch transmission.

The current model of itch spinal circuitry positions neurons that express the gastrin-releasing peptide receptor (GRPR) as a gate for itch input [37,38,45,61,62]. We explored the possibility that NK1R and GRPR neurons might represent an overlapping dorsal horn neuron subpopulation that modulates itch.

In this study, we demonstrate that activation of NK1R spinal neurons potentiates itch, while inhibition of NK1R spinal neurons attenuates itch. We highlight that, despite being widely used as a selective marker of spinal projection neurons, NK1R is expressed predominately in interneurons within the dorsal horn. Moreover, we reveal a previously unappreciated role for NK1R interneurons in spinal itch transmission, and that these NK1R neurons comprise a subpopulation of excitatory GRPR interneurons within the superficial dorsal horn.

## Materials and Methods

### Animals

All animals were cared for in compliance with the National Institutes of Health guidelines and approved by the University of Pittsburgh Institutional Animal Care and Use Committee.

Experiments were conducted on both male and female mice, with littermates randomly assigned to experimental groups. Adult, C57BL/6 mice were obtained from Charles River (strain 027) and allowed to acclimate to the University of Pittsburgh’s vivarium for at least one week prior to beginning experimentation. Previously generated *Tacr1*^*CreER*^ mice (also referred to as NK1R-CreER mice [27]) were bred in house and maintained on a C57BL/6 background. For histology and *in situ* hybridization studies, *Tacr1*^*CreER*^ mice were crossed with Cre-dependent *Rosa*^*tdT*^ reporter mice (Jackson Laboratory, stock no. 007909). For two-photon Ca^2+^ imaging studies, *Vglut2*^*Cre*^ mice (Jackson Laboratory, stock no. 016963) were crossed with Cre-dependent *Rosa*^*GCaMP6s*^ mice (Jackson Laboratory, strain no. 028866).

Mice were housed in cages of 4 (males) or 5 (females) in the animal facility under a 12 hr light/dark cycle (7AM – 7PM) and provided food and water *ad libitum*. Cages were lined with woodchip bedding. Behavioral studies on and tissue harvesting from C57BL/6 and *Tacr1*^*CreER*^ mice began when animals were 6-8 week old. Spinal cord Ca^2+^ imaging studies were similarly performed on 6-8 week old *Vglut2*^*Cre*^::*Rosa*^*GCaMP6s*^ mice.

### Tamoxifen administration

Tamoxifen (Sigma, T5648) was dissolved in corn oil (Sigma, C8267) by shaking it at 37°C to create a 20 mg/mL stock solution. This solution was stored, protected from light, at 4°C for up to 1 week. Tamoxifen was administered intraperitoneally at 75 mg/kg for 5 consecutive days.

*Tacr1*^*CreER*^;*Rosa*^*tdT*^ mice were injected with tamoxifen at ~2.5 weeks of age. Intraspinal viral injections were performed on ~ 4 week old *Tacr1*^*CreER*^ mice, and tamoxifen administration began 48 after surgery.

### Intraspinal viral injections

Mice were anesthetized with a ketamine/xylazine cocktail, 87.5 mg/kg/12.5 mg/kg, i.p.. Opthalmic ointment (Dechra, 12920060) was applied to the eyes. The back was shaved, and the local antiseptic betadine (Fischer Scientific, 19-061617) was applied to the skin. Using a scalpel, a skin incision was made over the T12-L3 vertebrae. Small scissors were used to cut through fascia, and #5 forceps were used to separate muscle from the T12-T13 vertebrae, exposing the intervertebral space above the left L3/L4 spinal segments [10,23]. Laminectomies were avoided to minimize spinal cord trauma. A glass capillary was carefully lowered down 300 μm from the surface of the dura, taking care to leave the posterior spinal artery intact. Once lowered, 500 nL of virus was injected using a NanoInject III Programmable Nanoliter Injector (Drummond Scientific Company) mounted on a stereotax (Kopf, Model 942) at a flow rate of 5 nL/sec, and the glass capillary was left in place for a total of 5 min and then slowly withdrawn. The skin incision was closed with 5-0 vicryl suture (Ethicon, VCP493G). Post-surgery, mice were injected subcutaneously with 0.3 mg/kg buprenorphine and 5 mg/kg ketoprofen and allowed to recover on a heating pad. On average, unilateral intraspinal viral injections resulted in 1.6 mm of rostral-caudal viral spread, spanning the L3-L5 spinal segments. Behavioral tests began 4 weeks after viral injection to allow for maximal and stable viral expression. At the conclusion of behavioral studies, intraspinal viral injections were confirmed with histology (described below). One Tacr1::hM3Dq-mCherry mouse was excluded from analyses because there was no hM3Dq-mCherry detected within the lumbar spinal segment.

### Viral constructs

For chemogenic and anterograde tracing studies, an adeno-associated virus (AAV2) coding for Cre-dependent hM3Dq-mCherry, AAV2-hSyn-DIO-hM3D(Gq)-mCherry (5.8 × 10^12^ particles/mL; Addgene, Catalog number 44361-AAV2, lot number V12832) was injected intraspinally; these animals are referred to as Tacr1::hM3Dq-mCherry mice throughout the text. AAV Cre-dependent fluorescent reporters: AAV2-EF1a-DIO-eYFP (4.6 × 10^12^ particles/mL; UNC Vector Core, Lot number AV4842E; 7 mice) or AAV2-hSyn-DIO-mCherry (4.7 × 10^12^ particles/mL; Addgene, Catalog number 50459-AAV2, lot number V27924; 3 mice) were used as controls. For simplicity, mice that received either control virus are referred to as Tacr1::control throughout the text. Viruses were stored in aliquots at −80°C until use.

### Chemogenetic studies

Based off of previous publications [30,47], mice were administered the water-soluble version of clozapine-N-oxide, clozapine N-oxide hydrochloride (CNO, Tocris, Cat No. 6329), which was dissolved in sterile saline and delivered at 5 mg/kg, i.p.. Behavioral testing was completed within 2.5 hr of CNO injection. Baseline nociceptive withdrawal testing (heat, cold, mechanical) took place at least 1 day prior to CNO administration.

### Behavioral studies

All behavioral studies were performed by the same tester to minimize experimenter variability. The experimenter was blind to all experimental conditions throughout data acquisition and analysis. Each behavioral study was performed on two different cohorts of mice for a total of 8-10 mice per experimental group across cohorts. For each cohort, the experimenter was blind to drug treatment or virus group until the completion of data analysis. Injection sites were shaved at least 24 hr prior to testing. On testing days, mice were acclimated to the testing apparatus in plexiglass boxes for at least 30 min prior to the start of testing. When applicable, once all animals in a cohort were injected, the experimenter left the room. Experiments were performed during the light cycle between 10AM and 5PM.

#### Intrathecal injections

Intrathecal injections were performed on awake, behaving animals to avoid effects of anesthesia on behavior as previously described, within minor modifications [33,44]. Mice were pinned down by the pelvic girdle, and drugs were delivered between the L5 and L6 vertebrae using a 30G needle attached to a 25 μL Hamilton syringe. Successful insertion of the needle into the intervertebral space was indicated by a sudden lateral movement of the tail. Thereafter, the drug was slowly injected at a rate of 1 nL/s, and the needle was held in place for an additional 5 s to minimize backflow. Only animals with successful injections were included in data analysis. The following drugs were delivered in a volume of 5 μL sterile saline: SP (400 ng, Sigma, S6883), selective NK1R agonist GR 73, 632 (40 ng, Tocris, 1669), gastrin-releasing peptide (GRP, 286 ng, Tocris, 1789), NK1R antagonist CP 99994 (18 μg, Tocris, 3417). Sterile saline was delivered as a vehicle control.

#### SP-, selective NK1R agonist-, and GRP-evoked behavior

Mice were injected intrathecally with SP, the selective NK1R agonist GR 73, 632, or vehicle as described above. Immediately following injection, animals were returned to their behavior boxes and their activity was video recorded to assess spontaneous behaviors. The three most prevalent behaviors elicited by SP and GR 73, 632 included scratch bouts, biting bouts, and head grooming. A scratch bout was defined as a rapid back and forth movement of the hindlimb (in this case, usually directed towards the abdomen) that ended in either licking/biting of the hindpaw or returning the hindpaw to the floor. A biting bout was defined as contact of the snout with the abdomen, ending with the mouse raising its head away from its body. Head grooming was defined as a single bilateral forepaw stroke, moving from the caudal to rostral end of the cheek, a stereotyped movement within mouse self-grooming chains [32]. The number of scratch bouts, biting bouts, and head grooming events were quantified for 5 min (SP) or 20 min (GR 73, 632) following intrathecal injection. In a separate experiment, mice were injected intrathecally with GRP and scratch bouts, biting bouts, and head grooming events were quantified for the subsequent 30 min.

#### Chloroquine-induced itch behavior

In pharmacological inhibition studies, chloroquine (100 μg in 5 μL sterile saline, Sigma, C6628) was administered intradermally into the calf 15 min following intrathecal injection of either CP 99994 or saline. Chemogenetic studies similarly used the calf model of itch [2,36] because intraspinal viral injections were localized to the lumbar spinal cord. Chloroquine (100 μg in 5 μL) was administered intradermally into the calf ipsilateral to intraspinal viral injection 120 min after the injection of CNO. For both studies, mice were video recorded for 30 min following the chloroquine injection, and the duration of time spent biting the calf was quantified off-line in 5-min intervals.

#### CNO-evoked spontaneous behavior

Immediately after administration of CNO, mice were video recorded for 60 min. The duration of all behaviors directed towards the hindlimb ipsilateral to the intraspinal viral injection, including hindpaw licking, lifting, flinching, as well as calf or haunch biting or grooming, was quantified off-line in 5-min intervals.

#### Heat sensitivity (Hargreaves test)

Animals were acclimated on a glass plate held at 30°C (IITC). Beginning 60 min following CNO administration, a radiant heat source (active intensity = 10%; intermittent intensity = 3%) was applied to the glass beneath the hindpaw and latency to paw withdrawal was recorded [24]. Three trials were conducted on each paw, with at least 5 min between testing the opposite paw and at least 10 min between testing the same paw. A cut off latency of 20 s was set to avoid tissue damage. Values across trials were averaged to determine withdrawal latency of each paw.

#### Cold sensitivity (Cold plantar assay)

Cold sensitivity was measured as previously described [15,16]. Mice were acclimated to a 1/4” glass plate and a cold probe was made by packing finely crushed dry ice into a modified 3 mL syringe 1 cm in diameter. Beginning 25 min after CNO administration, the cold probe was applied to the glass beneath the plantar surface of the hindpaw and the latency to paw withdrawal was recorded. Three trials were conducted on each hindpaw, with 5 min between trials on opposite paws, and 10 min between trials on the same paw. A cut off latency of 20 s was used to prevent tissue damage. Withdrawal latencies for each paw were determined by averaging values across trials.

#### Mechanical sensitivity (von Frey)

Mechanical sensitivity was measured using the simplified up-down (SUDO) method of the von Frey test [11]. Beginning 40 min following CNO administration, calibrated von Frey filaments (North Coast Medical Inc.) were applied to the plantar surface of the hindpaw for 2 s. For each trial, a total of 5 filament applications was conducted and used to calculate the paw withdrawal threshold. Three trials were conducted on each hindpaw, with 5 min between trials on opposite paws, and 10 min between trials on the same paw. The average paw withdrawal threshold from all three trials are reported in units of pressure (g/mm^2^) to account for differences in filament surface areas.

#### Capsaicin-induced nocifensive behavior

40 min after CNO injection, 10 μL of 0.001% capsaicin (dissolved in 10% ethanol, 0.5% Tween-80 in sterile saline; Sigma, M2028) was injected intraplantar into the hindpaw ipsilateral to the intraspinal viral injection and mice were video recorded for 15 min. The duration of nocifensive behaviors including lifting, licking, or shaking the hindpaw was quantified off-line in 5-min intervals.

### Stereotaxic injection and retrograde labeling

Mice were anesthetized with isoflurane (induction: 4%, maintenance: 2%) and head-fixed in a stereotaxic frame (Kopf, model 942). Ophthalmic ointment was applied to the eyes, the scalp was shaved, and local antiseptics (betadine and ethanol) were applied before using a scalpel to make a midline incision to expose the skull. The skull was leveled using cranial fissures as landmarks. To target the lateral parabrachial nucleus (lPBN), a drill bit (Stoelting, 514551) was used to create a burr hole at the following empirically derived coordinates: AP −5.11 mm, ML ±1.25 mm, DV −3.25mm. A glass capillary was carefully lowered through the burr hole to the injection site, and a Nanoinject III was used to deliver 500 nL of the fluorescently conjugated retrograde tracer cholera toxin subunit B-Alexa Fluor 647 (CTB, Thermofisher C34778, C22843) at a rate of 5 nL/s. The glass capillary was left in place for a total of 5 min and then slowly withdrawn. The scalp was sutured closed with 5-0 vicryl suture. Post-surgery, mice were injected subcutaneously with 0.3 mg/kg buprenorphine and 5 mg/kg ketoprofen and allowed to recover on a heating pad. Immunohistochemistry studies were conducted at least 10 days after CTB injection into the lPBN to allow for maximal retrograde labeling of spinal projection neurons.

### Immunohistochemistry

For histology studies, *Tacr1*^*CreER*^ mice expressing hM3Dq-mCherry (virally-mediated) and CTB-647 or tdTomato (constitutive reporter) were anesthetized with urethane and perfused with 4% paraformaldehyde. Lumbar spinal cord, dorsal root ganglia, and brain were dissected and post-fixed in 4% paraformaldehyde at 4°C for either 2 hr (spinal cord, dorsal root ganglia) or overnight (brain). Tissues were washed in PBS and cryoprotected in 30% sucrose prior to cryosectioning. When directly mounted onto slides, 40 μm transverse spinal cord and 20 μm dorsal root ganglia sections were collected. When stained as free-floating sections, 60 μm transverse spinal cord and 60 μm coronal brain sections were collected and stored in PBS with 0.01% sodium azide. Tissues were incubated in a blocking solution consisting of 10% normal donkey serum (Jackson ImmunoResearch, 017-000-121) and 0.3% Triton-X 100 solution (Sigma, 93443) in PBS for 1 hr at room temperature. The primary antibodies rabbit anti-RFP (Rockland, 600-401-379), guinea pig anti-cFOS (Synaptic Systems, 226 005), and rabbit anti-NK1R (Sigma, S8305, 1:10,000) were diluted in antibody buffer consisting of 5% normal donkey serum and 0.3% Triton-X 100 in PBS. Tissue was incubated in primary antibody overnight at 4°C, then washed in PBS three times for 10 min. The secondary antibodies donkey anti-rabbit Alexa-fluor 555 (Invitrogen, A-31572, 1:500) and goat anti-guinea pig Alexa-fluor 488 (Invitrogen, SA5-10094, 1:500), were diluted in antibody buffer and applied for 1 hr (mounted sections) or 2 hr (free-floating sections). Finally, tissues were washed again in PBS three times for 10 min and mounted with Prolong Gold with Dapi (Invitrogen, P36931).

In experiments evaluating expression of AAV2-hSyn-DIO-hM3D(Gq)-mCherry in the spinal cord and dorsal root ganglia, sections were mounted and stained directly on Super Frost Plus slides (Fisher Scientific, 12-550-15). In studies visualizing NK1R immunoreactivity, as well as the brain targets of *Tacr1*^*CreER*^ spinal projection neurons, sections were stained free-floating. In experiments evaluating the percentage of *Tacr1*^*CreER*^ spinoparabrachial neurons versus interneurons, only mice with confirmed on-target CTB injections into the PBN were analyzed. CTB-647 and tdTomato signals were not amplified.

### cFOS induction

To verify that systemic CNO administration caused hM3Dq – and in turn neuronal – activation, Tacr1::hM3Dq-mCherry and Tacr1::control mice were injected with 5 mg/kg CNO i.p. 90 min prior to perfusion with 4% paraformaldehyde, in line with the time course of behavioral studies. Spinal cord tissue was processed and immunostained for as described above.

### Fluorescence in situ hybridization (RNAscope)

Mice were anesthetized with isoflurane and rapidly decapitated. The L3/L4 spinal cord segments were quickly removed within 5 min, placed into OCT, and flash frozen using 2-methylbutane chilled on dry ice. Tissue was kept on dry ice until cryosectioning. 15 μm cryosections were mounted directly onto Super Frost Plus slides, and fluorescence *in situ* hybridization (FISH) studies were performed according to the protocol for fresh-frozen samples using the RNAscope Multiplex Fluorescent v2 Assay (Advanced Cell Diagnostics, 323100) with minor modifications. Briefly, spinal cord tissue was dehydrated with ethanol and fixed for 15 min in ice-cold 4% PFA. Sections were treated with hydrogen peroxide for 10 min at room temperature, followed by Protease IV for 15 min at room temperature. Probes for *Tacr1* (Advanced Cell Diagnostics, Mm-Tacr1, 428781)*, tdTomato* (-tdTomato, 317041), *Grpr* (Mm-Grpr, 517631), and *Slc17a6* (referred to here as *Vglut2*, Mm-Slc16a6, 319171) were hybridized for 2 hr at 40°C in a humidified oven, then stored overnight in 5X saline sodium citrate. After rinsing in wash buffer, a series of incubations was then performed to amplify and develop hybridized probe signal using TSA Plus Fluorophores (Perkin Elmer, NEL741001KT and NEL744E001KT). Slides were mounted with Prolong Gold with DAPI.

### Imaging and Quantification

Sections were imaged at full tissue-thickness using an upright epifluorescent microscope (Olympus BX53 with UPlanSApo 4x, 10x, or 20x objectives) or a confocal microscope (Nikon A1R with 20x or 60x oil-immersion objectives). All analysis was completed off-line using FIJI software (Image J, NIH). For FISH experiment quantification, 3-4 spinal cord sections/mouse were manually quantified from 4-5 mice/experiment. A positive cell was defined as a cell with a clearly defined nucleus and fluorescent signal forming a ring around the nucleus. To quantify the number of recombined Tacr1::hM3Dq-mcherry neurons from the onset to offset of viral spread, neurons from every 6^th^ section (thus, every 240 μm) were quantified and their laminar distribution was noted (n=7 mice). For CTB-backlabeled and cFOS neuron quantification, 4-5 sections/mouse were manually counted from 3-4 mice/condition.

In analyses of the spatial distribution of *Tacr1*, Tacr1::hM3Dq-mCherry, and CTB neurons, the superficial dorsal horn (SDH) was defined as the region 65 μm below the surface of the dorsal horn gray matter, roughly corresponding to laminae I and II_o_. This distance was determined by measuring the distance from the surface of the dorsal horn gray matter to the bottom of the *Grpr* band in FISH sections, based on previous studies demonstrating that GRPR neurons are restricted to the superficial dorsal horn [9,61,62]. The deeper dorsal horn (DDH) consisted of the remaining lamina, roughly III-VI of the dorsal horn. The lateral spinal nucleus (LSN) neurons were defined as those located lateral to the dorsal horn gray matter, within the dorsolateral funiculus.

### Two-photon Ca^2+^ imaging

As described previously [22], the C2 – S6 spinal cord segments were dissected from *Vglut2*^*Cre*^::*Rosa*^*GCaMP6s*^ mice and placed in a Sylgard recording chamber designed for pharmacology providing fast and uniform fluid exchange. Using a Leica SP-5 multiphoton microscope coupled to a Chameleon Ultra tunable Ti:Sapphire laser and a Leica 20x (NA 1.00), the lumbar (L1-3) superficial grey matter (0 - 100 μm, encompassing laminae I and II) was imaged at 3-4 Z-planes at 1 Hz, sampling up to ~200 excitatory neurons in a given experiment within a typical ~0.13 mm^2^ field of view. Throughout imaging experiments, the recording chamber was continuously perfused with normal aCSF solution (in mM: 117 NaCl, 3.6 KCl, 2.5 CaCl_2_, 1.2 MgCl_2_, 1.2 NaH_2_PO_4_, 25 NaHCO_3_, 11 glucose) saturated with 95% O_2_ and 5% CO_2_ at 30°C. TTX (500 nM, Tocris 1069) was added to the bath in order to identify neurons that responded directly to SP (1μM, Sigma, S6883) or GRP (300 nM, Tocris, 1789), which were applied 5 min apart, in random order. Agonist concentrations were selected based on previous *in vitro* spinal cord physiology studies [6,38,41,42,45]. At the conclusion of experiments, modified aCSF containing 30 mM K^+^ was perfused on the spinal cord to activate and visualize all GCaMP6-expressing neurons, as well as to assess their viability.

For image processing, a Suite2p pipeline, custom-tuned for dorsal horn neurons was used for image registration and signal extraction. ImageJ was used for all other image processing. Responders were defined as neurons that had both a peak value in ΔF/F of ≥ 80% and a ΔF/F of ≥ 4 standard deviations calculated from the 30 seconds prior to drug perfusion. Within GRP responsive cells, Ca^2+^ oscillations, likely arising from intracellular stores [34], were noted and if an individual neuron’s oscillations extended into the period of time during SP application, the cell was excluded from analysis.

### Statistical Analyses

Microscope Excel, GraphPad Prism, and the R treemap package were used for data organization and statistical analyses. Statistical significance was indicated by p ≤ 0.05 determined using Student’s t-Test, Mann-Whitney *U* Test, two-way RM ANOVA, two-way ANOVA, or a mixed-effects model. The Holm-Sidak test was used to correct for multiple comparisons, when applicable. See figure legends for experiment-specific details. All values are presented as mean ± SEM.

## Results

### NK1R spinal neurons bidirectionally modulate itch-related behaviors

When injected into the skin, SP can elicit itch-related behaviors such as scratching [7,8]. We tested whether SP similarly acts as a pruritogen within the spinal cord. C57BL/6 mice were injected intrathecally with SP or saline, and their spontaneous behaviors were quantified (Figure 1a). Compared to control mice injected with saline, mice injected with SP exhibited significantly more spontaneous itch-related behaviors, including scratch bouts, biting bouts, and head grooming events (Figure 1b). The effect of SP was immediate and subsided within 5 min, likely due to rapid internalization of NK1R upon binding of SP [39,67].

**Figure 1.**
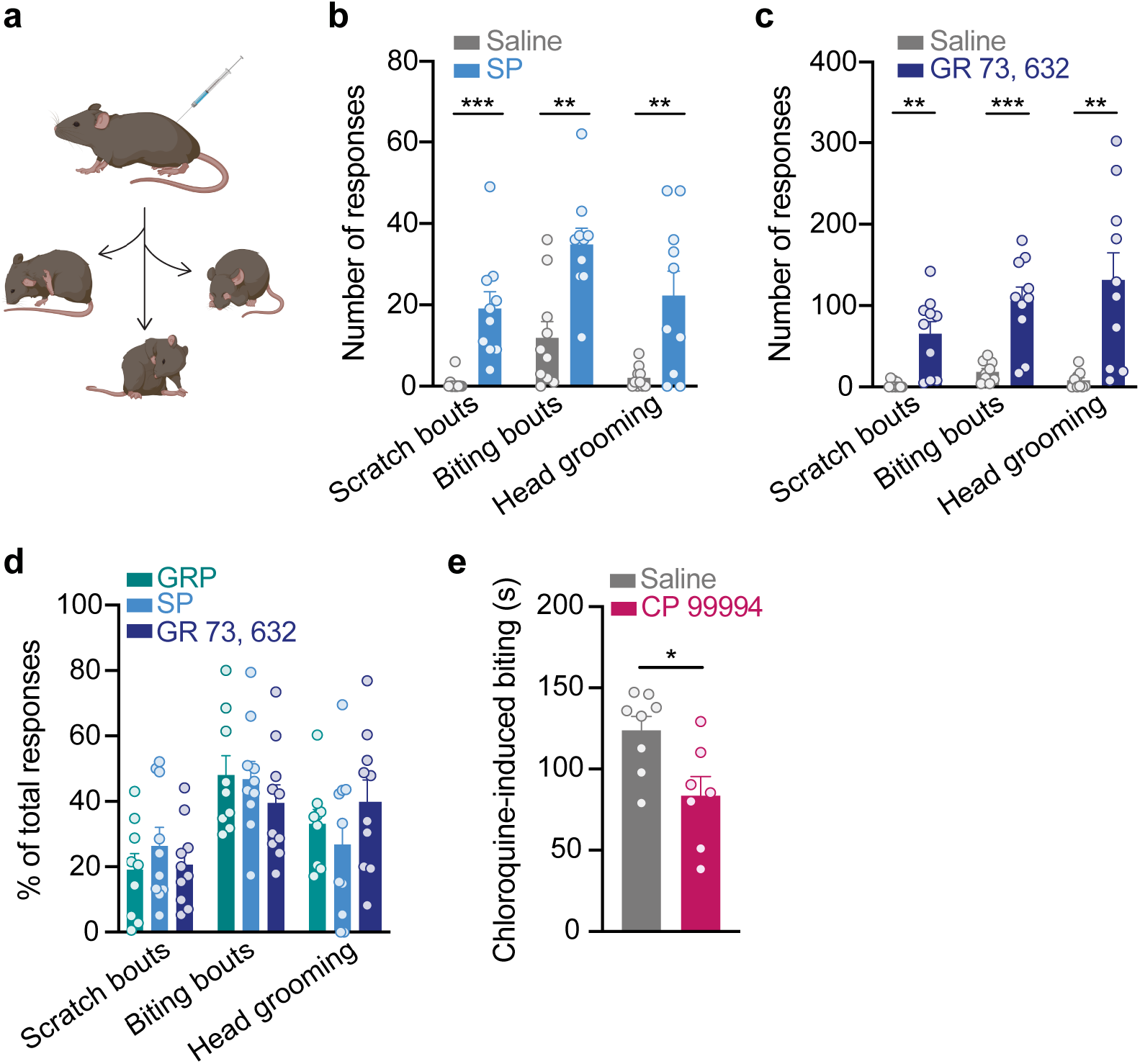
Spinal NK1R bidirectionally modulates itch. **a**, C57BL/6 mice were injected intrathecally with either saline, SP (400 ng), the selective NK1R agonist GR 73, 632 (40 ng), or GRP (285 ng), and three spontaneous itch-related behaviors were quantified: scratch bouts, biting bouts, and head grooming events. **b**, Intrathecal SP elicited significantly more scratch bouts, biting bouts, and head grooming responses than saline in the 5 min immediately following injection (Student’s t-Test, Holm-Sidak correction for multiple comparisons, scratch bouts: \*\*\**p*=9.52 × 10^−4^ , *t*=4.44, df=18, biting bouts: \*\**p*=1.60 × 10^−3^ , *t*=4.02, df=18, head grooming: \*\**p*=3.58 × 10^−3^ , *t*=3.35, df=18; n=10 mice/group). **c**, Intrathecal GR 73, 632 significantly increased the number of scratch bouts, biting bouts, and head grooming responses compared to saline in the 20 min immediately following injection (Student’s t-Test, Holm-Sidak correction for multiple comparisons, scratch bouts: \*\**p*=2.01 × 10^−3^, *t*=3.96, df=17, biting bouts: \*\*\**p*=5.73 × 10^−4^, *t*=4.74, df=17, head grooming: \*\**p*= 2.66× 10^−3^, *t*=3.51, df=17; n=9-10 mice/group). **d**, The percentage of scratch bouts, biting bouts, and head grooming responses out of total responses to intrathecal injection of SP and GR 73, 632 were equivalent to that elicited by the well-established pruritogen, GRP (2-way ANOVA, behavioral response × agonist, *p*=0.296, *F*(4,78)=1.25, n=9-10 mice/group). Behavioral responses to GRP were quantified for the 30 min immediately following injection. **e**, Intrathecal pretreatment with the NK1R antagonist CP 99994 (18 μg) significantly decreased the duration of site-directed biting in response to an intradermal injection of chloroquine into the calf (100 μg/5μL) compared to saline (Student’s t-Test, \**p*=0.016, *t*=2.77, df=13; n=7-8 mice). SP, substance P; GRP, gastrin-releasing peptide. Data are shown as mean ± SEM, with open circles representing individual mice.

Because SP can bind at lower affinities to other tachykinin receptors that are expressed on spinal neurons [57] – NK2R and NK3R – we tested whether selective activation of NK1R similarly elicited spontaneous itch-related behaviors. Indeed, like SP, intrathecal injection of the selective NK1R agonist GR 73, 632 caused a significant increase in the number of scratch bouts, biting bouts, and head grooming responses relative to saline (Figure 1c). In contrast to SP, the effects of GR 73, 632 were longer lasting, and continued for at least 20 min, suggesting that NK1R internalization does not occur as rapidly upon binding of GR 73, 632.

These findings are in agreement with previous reports that intrathecal administration of SP elicits scratching and biting behavior in rodents [29,51,55]. In some cases, SP-induced scratching and biting have been referred to as pain-related behaviors. However, the behavioral responses to intrathecal injection of SP and GR 73, 632 closely resembled the those to intrathecal injection of the well-established pruritogen, gastrin-releasing peptide (GRP). Comparing mice that were injected intrathecally with either GRP or NK1R agonists, we determined the percentage of total behavioral responses represented by scratch bouts, biting bouts, and head grooming for each agonist. Notably, there was no difference in the percentage of scratch bouts, biting bouts, and head grooming events elicited by SP, GR, 73, 632, or GRP (Figure 1d). Thus, we favor the idea that like GRP, behaviors triggered by spinal NK1R activation are itch-related. Increased head grooming is an interesting observation in these experiments because it is not a canonical itch-related behavior. It might, however, represent an indirect response to itch. There is growing evidence that itch is aversive and increases anxiety in rodent models [53,54], and increased anxiety is in turn associated with heightened rodent self-grooming [31,32].

Since activation of spinal NK1R potentiated itch, we then asked whether inhibition of spinal NK1R blocked itch. We intrathecally delivered the NK1R antagonist CP 99994 at a dose previously shown to inhibit pain behaviors [12], then injected the pruritogen chloroquine intradermally into the calf to elicit itch. When pruritogens are injected into the calf, mice bite the injection site [36]. As expected, compared to mice pretreated with saline, mice pretreated with CP 99994 exhibited significantly less calf-direct biting in response to chloroquine (Figure 1e), complementing previous findings [1,3,4,19]. Taken together, these results indicate that NK1R spinal neurons mediate itch behaviors.

### *Tacr1* is broadly expressed within the spinal cord dorsal horn

After establishing that NK1R spinal neurons drive itch-related behaviors, our overarching goal was to understand which NK1R neurons within the spinal cord contribute to itch. As a first step, we sought to characterize the laminar distribution of NK1R neurons within the spinal cord dorsal horn. We performed fluorescence in situ hybridization (FISH) for *Tacr1*, the gene encoding NK1R, and quantified the number of *Tacr1*-expressing neurons located within the superficial dorsal horn (SDH), deeper dorsal horn (DDH), and lateral spinal nucleus (LSN) (Figure 2a,b).

**Figure 2.**
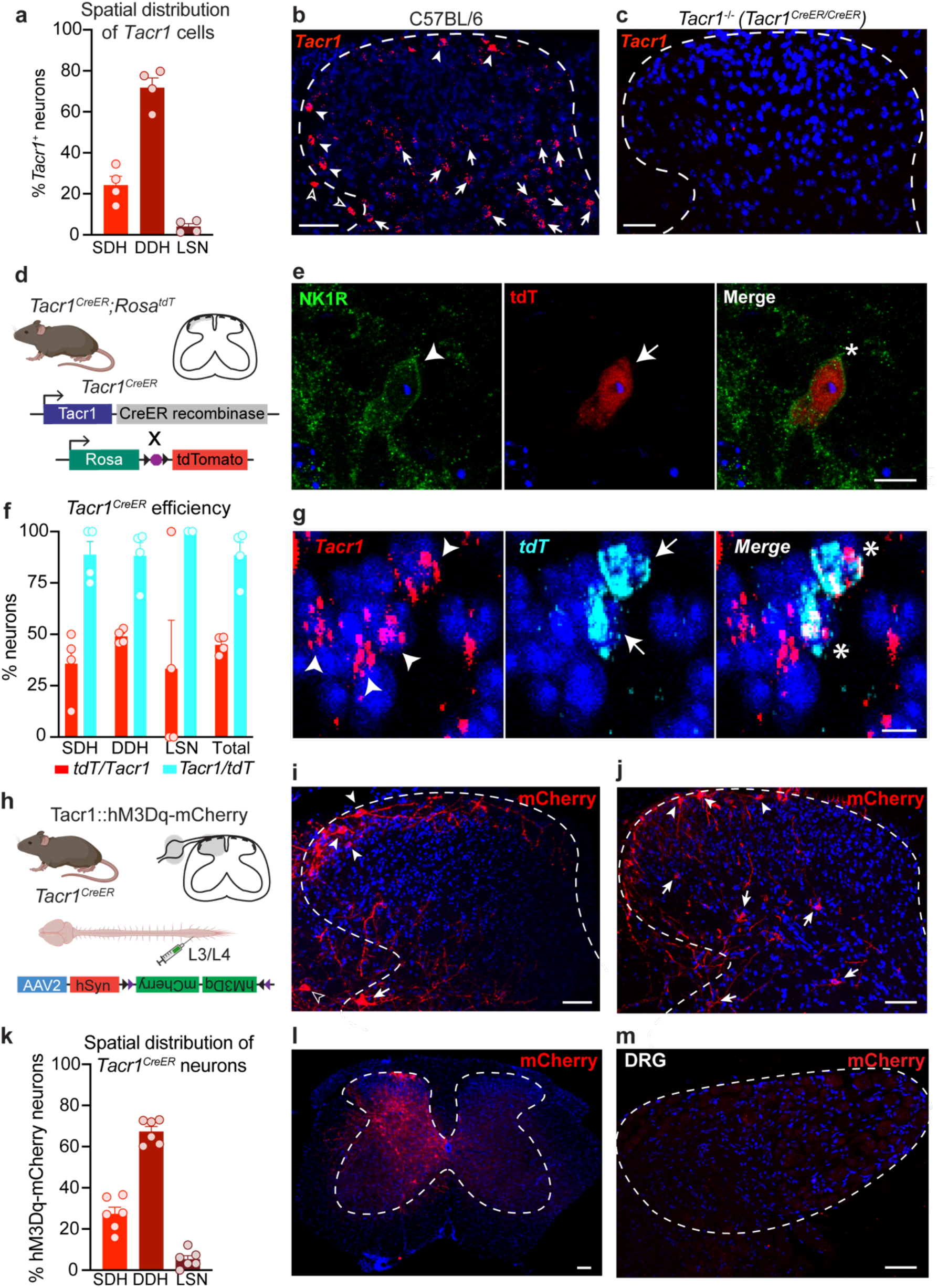
Characterization of *Tacr1* expression in C57BL/6 and *Tacr1*^*CreER*^ mice. **a**, Percentage of *Tacr1* neurons located within the SDH, DDH, and LSN of the lumbar spinal cord (n=4 mice), visualized via **b**, FISH in C57BL/6 mice. Filled arrowheads, SDH; filled arrows, DDH; empty arrowheads, LSN. **c**, No *Tacr1* signal was detected by FISH in the lumbar spinal cord of *Tacr1*^*CreER/CreER*^ mice, which are equivalent to NK1R^−/−^ mice. Scale bars in **b, c**, 50 μm. **d**, To evaluate the efficiency and specificity of *Tacr1*^*CreER*^-mediated recombination within the lumbar spinal cord, *Tacr1*^*CreER*^;*Rosa*^*tdT*^ mice were generated. **e**, Representative IHC image showing colocalization of NK1R immunoreactivity (green) and *Tacr1*^*CreER*^-mediated tdT expression (red) in an SDH neuron. Scale bar, 10 μm. **f**, Quantification of the colocalization of *Tacr1* and *tdT* mRNA within SDH, DDH, and LSN neurons (n=4 mice). **g**, Representative FISH image showing *Tacr1*^*CreER*^-mediated tdT mRNA expression (cyan) in *Tacr1*-expressing (red) spinal dorsal horn neurons. Scale bar, 10 μm. **h**, *Tacr1*^*CreER*^ mice received intraspinal injections of the Cre-dependent virus AAV2-hSyn-DIO-hM3Dq-mCherry to determine whether Cre-mediated recombination recapitulated the spatial distribution of endogenous *Tacr1* expression. **i, j**, Representative IHC images of the spatial distribution of hM3Dq-mCherry neurons (red) throughout the spinal cord dorsal horn. Filled arrowheads, SDH; filled arrows, DDH; empty arrowheads, LSN. Scale bars, 50 μm. **k**, Percentage of hM3Dq-mCherry neurons located within the SDH, DDH, and LSN of the lumbar spinal cord (n=6 mice). **l**, Representative IHC image of the lumbar spinal cord showing that viral-vector mediated Cre-dependent recombination in *Tacr1*^*CreER*^ mice, indicated by mCherry expression (red), was largely restricted to the side ipsilateral to the intraspinal viral injection. Scale bar, 50 μm. **m**, Representative IHC image of the corresponding ipsilateral dorsal root ganglia (DRG), which lacks mCherry (red), indicating that viral vector-mediated Cre-dependent recombination was restricted to the spinal cord of *Tacr1*^*CreER*^ mice. Scale bar, 50 μm. FISH, fluorescence in situ hybridization; SDH, superficial dorsal horn; DDH, deeper dorsal horn; LSN, lateral spinal nucleus. Data in **a, f, k** are shown as mean ± SEM, with open circles representing individual mice.

*Tacr1* neurons were spatially distributed throughout the dorsal horn. However, only a fraction of *Tacr1* neurons (24.2% ± 4.3) were localized to the SDH, with the majority of *Tacr1* neurons (71.8% ± 4.8) instead found in the DDH. A small percentage of *Tacr1* neurons were located in the LSN (4.0% ± 1.5). Importantly, FISH was also performed on spinal cord tissue from a homozygous *Tacr1*^*CreER*^ mouse, which is homozygous null (NK1R^−/−^), and we detected negligible signal, confirming the specify of the *Tacr1* probe (Figure 2c).

Our findings demonstrate that *Tacr1* is broadly expressed throughout the superficial and deeper dorsal horn. However, NK1R has long been used as a marker for ascending spinal projection neurons that reside within laminae I and III-V of the dorsal horn, as well as the lateral spinal nucleus [18,50,56,65]. Spinal projection neurons of the anterolateral tracts are an extremely sparse population, comprising no more than 5% and 0.1% of total neurons within the SDH and DDH, respectively [64], but the total numbers of *Tacr1* neurons in our study suggest more widespread expression. Thus, these results suggest that *Tacr1* expression is not limited to spinal projection neurons, consistent with previous reports [21,63].

### *Tacr1*^*CreER*^ captures a subpopulation of *Tacr1* spinal dorsal horn neurons

Dissection of the role of NK1R neurons in spinal processing of itch has been limited by a lack of genetic tools that selectively target NK1R neurons. To gain genetic access to *Tacr1* neurons, we previously generated a *Tacr1*^*CreER*^ knock-in mouse line [27]. However, the efficiency and specificity of *Tacr1*^*CreER*^-mediated recombination within spinal neurons was unknown.

To assess the efficiency and specificity of our Cre line, we crossed *Tacr1*^*CreER*^ mice to mice harboring a Cre-dependent tdTomato (tdT) fluorescent reporter expressed under control of the *Rosa* locus (*Tacr1*^*CreER*^;*Rosa*^*tdT*^) (Figure 2d). Immunohistochemistry and FISH revealed tdT colocalization with NK1R at the protein and mRNA transcript level, respectively (Figure 2e,g). Quantification of *Tacr1* and *tdT* neurons in dual FISH experiments indicated that while Cre-mediated recombination is highly specific to *Tacr1* neurons (88.5% ± 6.3 of all *tdT* neurons expressed *Tacr1*), it only occurs in roughly half of all *Tacr1*-expressing cells (44.9% ± 2.2 of all *Tacr1* cells expressed *tdT*) (Figure 2f) across dorsal horn spatial regions.

Dissecting the functional role of NK1R spinal neurons in itch required selective targeting of *Tacr1*^*CreER*^ neurons via local injection of Cre-dependent viruses into the spinal cord. As a first step, we asked whether virally-mediated Cre-dependent recombination in the *Tacr1*^*CreER*^ mouse line recapitulated the spatial distribution of endogenous *Tacr1* expression. A Cre-dependent virus encoding a neuron-specific excitatory DREADD fused to an mCherry fluorescent reporter (Tacr1::hM3Dq-mCherry; AAV2-hSyn-DIO-hM3Dq-mCherry) was injected into the L3/L4 spinal segment of *Tacr1*^*CreER*^ mice (Figure 2h). We selected a Cre-dependent virus encoding a membrane bound receptor over a cytosolic reporter to allow for better visualization of hM3Dq-mCherry neuron processes. Lumbar spinal cord segments were collected from Tacr1::hM3Dq-mCherry mice and recombined spinal neurons were visualized using an anti-RFP antibody (Figure 2i,j). The spatial distribution of hM3Dq-mCherry neurons closely resembled that of *Tacr1* cells, with 27.3% ± 3.3 located in the SDH, 67.3% ± 2.4 in the DDH, and 5.4% ± 1.7 in the LSN (Figure 2k). Thus, *Tacr1*^*CreER*^ captures a subpopulation of *Tacr1* spinal dorsal horn neurons that is representative of the spatial distribution of all *Tacr1* neurons.

Notably, viral-vector mediated Cre-dependent recombination was largely restricted to the side ipsilateral to the intraspinal viral injection (Figure 2l). Moreover, viral transduction in *Tacr1*^*CreER*^ mice was restricted to the spinal cord, as hM3Dq-mCherry was not detected in the corresponding lumbar dorsal root ganglia (Figure 2m), in agreement with previous evidence that the AAV2 viral serotype is not retrogradely transported along axons [17].

### Chemogenetic activation of *Tacr1*^*CreER*^ spinal neurons preferentially increases itch

Next, to complement our pharmacology studies, we tested whether chemogenetic activation of *Tacr1*^*CreER*^ spinal neurons similarly increased itch-related behaviors. To selectively activate NK1R spinal neurons, Cre-dependent viruses encoding a neuron-specific excitatory DREADD fused to a fluorescent reporter (Tacr1::hM3Dq-mCherry; AAV2-hSyn-DIO-hM3Dq-mCherry) or a Cre-dependent control virus encoding only a fluorescent reporter (Tacr1::control; AAV2-hSyn-DIO-mCherry or AAV2-EF1a-EYFP) were injected into the L3/L4 spinal segment of *Tacr1*^*CreER*^ mice (Figure 3a). In Tacr1::hM3Dq-mCherry mice, systemic administration of CNO significantly increased site-directed biting in response to an intradermal injection of chloroquine into the calf compared to Tacr1::control mice (Figure 3b,c). The duration of cumulative chloroquine-induced calf biting in Tacr1::hM3Dq-mCherry mice was significantly correlated to the number of hM3Dq-mCherry neurons (Figure 3d). Thus, a significant amount of the variability in the chloroquine-induced biting of Tacr1::hM3Dq-mCherry mice is explained by variability in viral transduction.

**Figure 3.**
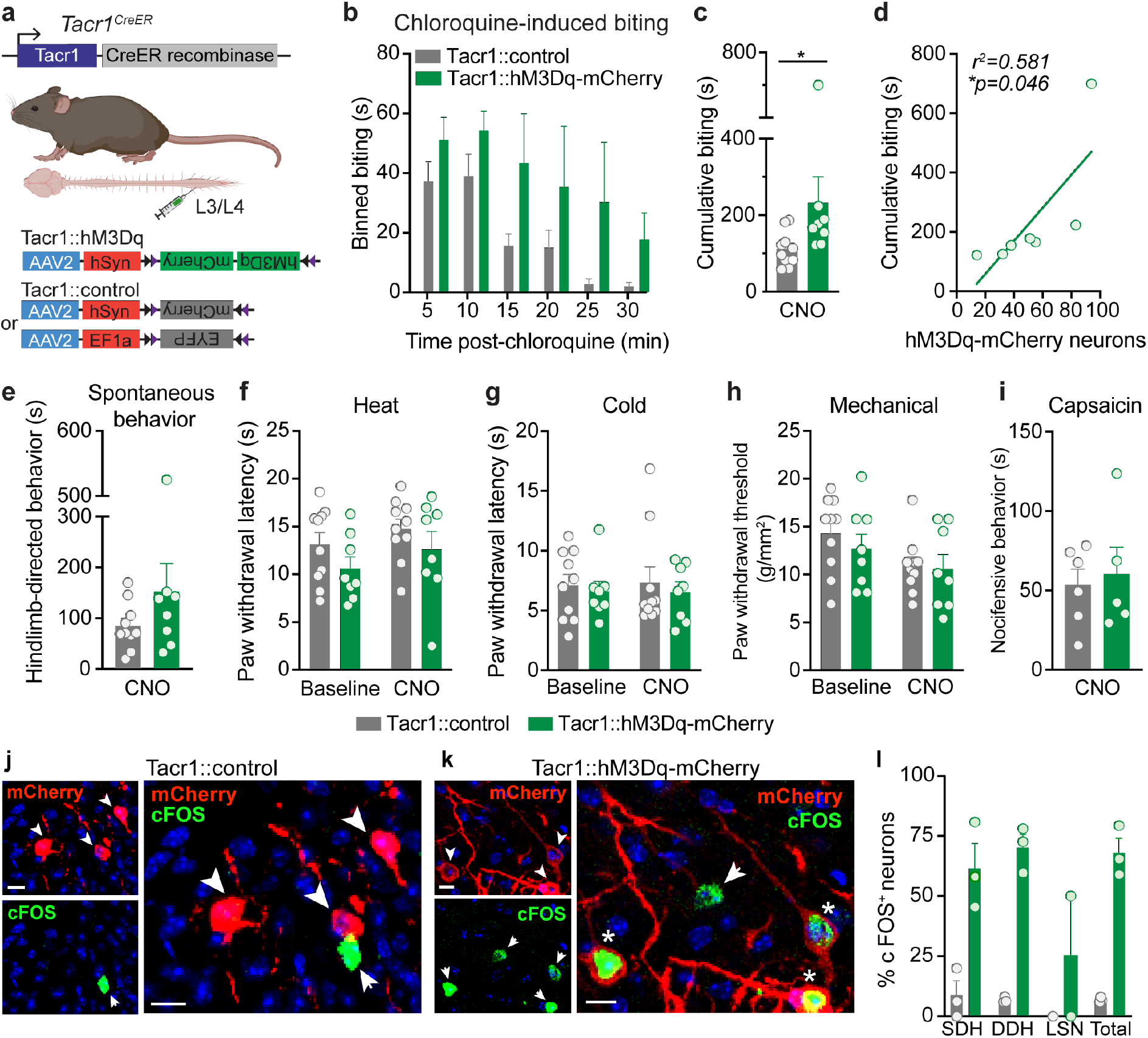
Chemogenetic activation of *Tacr1*^*CreER*^ spinal neurons preferentially increases itch. **a**, Strategy for selectively targeting *Tacr1*^*CreER*^ spinal neurons. To gain chemogenetic access to *Tacr1*^*CreER*^ spinal neurons, unilateral intraspinal viral injections of AAV2-hSyn-DIO-hM3Dq-mCherry (Tacr1::hM3Dq-mCherry) and either AAV2-hSyn-DIO-mCherry or AAV2-EF1a-DIO-EYFP (Tacr1::control) were targeted to the lumbar dorsal horn of *Tacr1*^*CreER*^ mice. **b**, Duration of site-directed biting in response to an intradermal chloroquine injection (100 μg/5 μL) into the calf following systemic CNO administration (5 mg/kg, i.p) in Tacr1::control and Tacr1::hM3Dq-mCherry mice, binned over time (n=8-10 mice/group). **c**, CNO administration significantly increased the cumulative duration of calf biting of Tacr1::hM3Dq-mCherry mice compared to Tacr1::controls (Mann-Whitney *U* test, \**p*=0.0155, *U*=13, df=16, n=8-10 mice/group). **d**, The duration of cumulative chloroquine-induced biting in Tacr1::hM3Dq-mCherry mice was significantly correlated with the number of recombined hM3Dq-mCherry neurons (Pearson’s Correlation, \**p*=0.0463, *r*^*2*^=0.581, *r*=0.762, n=7 mice). **e**, Chemogenetic activation of *Tacr1*^*CreER*^ neurons with CNO did not affect hindlimb-directed pain or itch spontaneous behaviors of Tacr1::hM3Dq-mCherry mice compared to Tacr1::controls (Mann-Whitney *U* test, *p*=0.315, *U*=28, df=16, n=8-10 mice/group). Hindpaw nociceptive withdrawal thresholds to **f**, heat (2-way RM ANOVA, CNO × DREADD, *p*=0.889, *F* (1,16) =0.0202, n=8-10 mice/group), **g**, cold (2-way RM ANOVA, CNO × DREADD, *p*=0.886, *F* (1,16)=0.0213, n=8-10 mice/group), and **h**, mechanical (2-way RM ANOVA, CNO × DREADD, *p*=0.520, *F* (1,16)=0.432, n=8-10 mice/group) stimuli were unchanged following CNO administration in Tacr1::hM3Dq-mCherry mice compared to Tacr1::controls. **i**, CNO administration did not alter the duration of nocifensive behaviors in response to intraplantar capsaicin (0.001% capsaicin, 10 μL) in Tacr1::hM3Dq-mCherry mice relative to Tacr1::controls (Student’s t-Test, *p*=0.729, *t*=0.358, *df*=9, n=5-6 mice/group). Representative IHC images of lumbar spinal cord sections demonstrating mCherry (red) and cFOS (green) expression in **j**, Tacr1::control and **k**, Tacr1::hM3Dq-mCherry mice 90 min following CNO administration, coinciding with the timeframe of behavioral testing. Scale bars, 10 μm. **l**, Quantification of cFOS+ mCherry neurons shows that CNO induced significantly more cFOS expression in Tacr1::hM3Dq-mCherry neurons compared to Tacr1::neurons (Mixed effects models, virus × spatial distribution, \*\**p*=0.0072, n=3 mice/group). **b,c, e-i, l**, Data are shown as mean ± SEM, with open circles representing individual mice.

Given the long-standing evidence that NK1R neurons are required for pain behaviors following injury [40,43,66,68], we next tested whether chemogenetic activation of *Tacr1*^*CreER*^ spinal neurons also increased nociceptive or acute pain-related behaviors. CNO administration did not produce spontaneous hindlimb-directed pain or itch behaviors in Tacr1::hM3Dq-mCherry mice compared to Tacr1::control mice (Figure 3e). Similarly, activation of *Tacr1*^*CreER*^ spinal neurons did not alter nociceptive withdrawal thresholds to heat, cold, or mechanical stimuli (Figure 3f-h) compared to controls, as measured by the Hargreaves test, cold plantar assay, and von Frey test, respectively. These findings are consistent with previous reports that nociceptive withdrawal thresholds are unchanged by neurotoxic ablation of NK1R in the absence of injury [40,43].

As a control for the possibility that the presence of the DREADD alone could influence baseline nociceptive withdrawal thresholds [52], baseline thresholds of Tacr1::control and Tacr1::hM3Dq mice were also measured in the absence of CNO. Expression of hM3Dq-mCherry alone did not affect baseline sensitivity to heat, cold, or mechanical stimuli compared to expression of control viruses (Figure 3f-h, Supp Table 1). Moreover, CNO did not have a frank effect on nociception, as administration did not alter the withdrawal thresholds of the hindpaw contralateral to intraspinal viral injection in Tacr1::Control or Tacr1::hM3Dq mice relative to baseline (Supp Figure 1a-c).

Lastly, using the intraplantar capsaicin model of neurogenic inflammation, we found that nocifensive behavioral responses such as licking or lifting of the hindpaw were unaffected by activation of *Tacr1*^*CreER*^ spinal neurons (Figure 3i). These findings diverge from previous reports that NK1R-expressing neurons are required for capsaicin-induced nocifensive behaviors [40,43,66].

To confirm that the dose of CNO used in behavioral studies was sufficient to activate Tacr1::hM3Dq-mCherry neurons, we evaluated cFOS expression as a surrogate marker for neuronal activation across the spinal cord dorsal horn (Figure 3j,k). CNO-induced cFOS expression was significantly higher in Tacr1::hM3Dq-mCherry neurons (67.9% ± 6.0 of total neurons) as compared to Tacr1::control neurons (6.9% ± 0.6 of total neurons) (Figure 3l).

Collectively, these findings suggest that chemogenetic activation of *Tacr1*^*CreER*^ spinal neurons preferentially increases itch behavior.

### *Tacr1*^*CreER*^ captures predominantly interneurons

Previous neurotoxic ablation studies were interpreted to support a role for NK1R spinal projection neurons in itch. However, data here suggest that ablation strategies would have likely targeted interneurons as well. To investigate this, we asked whether virally-captured neurons in Tacr1::hM3Dq-mCherry mice were spinal projection neurons or interneurons.

First, we leveraged AAV2-hSyn-DIO-hM3Dq-mCherry as an anterograde tracer and evaluated the brains of Tacr1::hM3Dq-mCherry mice for the central targets of ascending spinal projection neurons (Figure 4a). In line with previous findings in rodents [5,18], the most robust hM3Dq-mCherry processes were observed in the contralateral lateral parabrachial nucleus (lPBN), as well as the contralateral lateral and ventrolateral periaqueductal gray (PAG) (Figure 4b,c).

**Figure 4.**
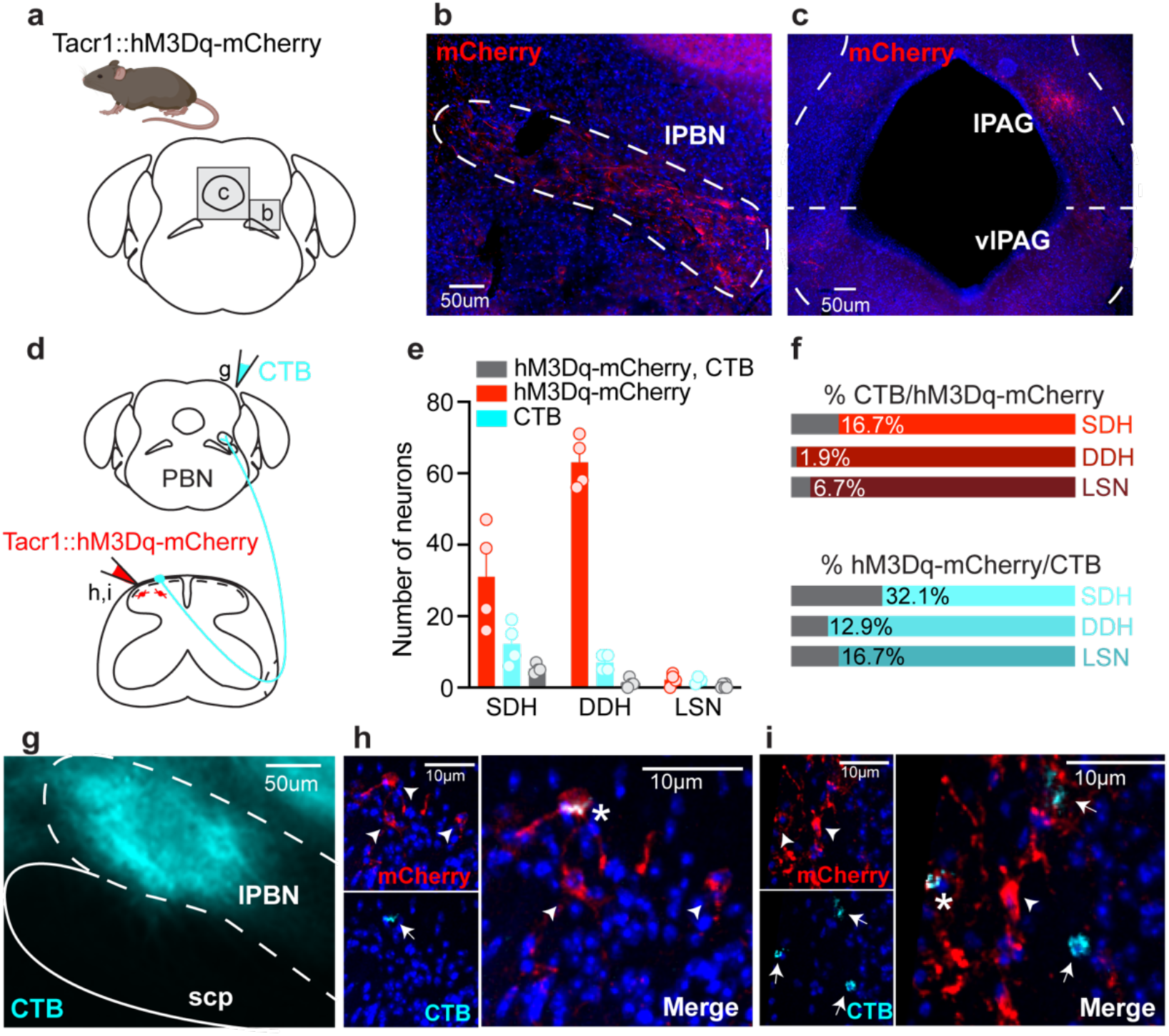
*Tacr1*^*CreER*^ spinal neurons are predominately interneurons. **a**, The brains of Tacr1::hM3Dq-mCherry neurons were collected and sectioned coronally to evaluate whether *Tacr1*^*CreER*^ captured spinal projection neurons. Representative immunostained coronal brain sections showing hM3Dq-mCherry processes (red) within the **b**, lPBN, as well as the **c**, lPAG and vlPAG contralateral to intraspinal viral injection. Scale bars, 50 μm. **d**, Experimental strategy to determine the relative proportion of Tacr1::hM3Dq-mCherry spinal projection neurons versus interneurons. The retrograde tracer CTB was injected into the contralateral lPBN of Tacr1::hM3Dq-mcherry mice to distinguish spinoparabrachial neurons (CTB and virally-mediated hM3Dq-mCherry expression) from interneurons (virally-mediated hM3Dq-mCherry expression only). **e**, Quantification of the number of neurons labeled by hM3Dq-mCherry, CTB, or both within the SDH, DDH, and LSN of Tacr1::hM3Dq-mCherry mice, revealing that the majority of *Tacr1*^*CreER*^ spinal neurons are local interneurons (n=4 mice). Data are shown as mean ± SEM, with open circles representing individual mice. **f**, Percentage of (top) total hM3Dq-mCherry neurons that were dual-labeled with CTB and (bottom) total CTB neurons that were dual-labeled with hM3Dq-mCherry within the SDH, DDH, and LSN, based off of number of neurons presented in **e**. **g**, Representative image of a targeted CTB injection (cyan) into the lPBN. Scale bar, 50 μm. **h, i**, Representative IHC images of lumbar spinal cord sections demonstrating the small extent of colocalization between hM3Dq-mcherry (red) and CTB (cyan). Scale bars, 10 μm. lPBN, lateral parabrachial nucleus; scp, superior cerebellar peduncle; lPAG, lateral periaqueductal gray; vlPAG, ventrolateral periaqueductal gray. SDH, superficial dorsal horn; DDH, deeper dorsal horn; LSN, lateral spinal nucleus.

Next, we performed dual-labeling studies to distinguish Tacr1::hM3Dq-mCherry spinal projection neurons from interneurons. Though anterograde tracing revealed hM3Dq-mCherry neuron processes in both the lPBN and the PAG, we chose to back label neurons from the lPBN because it has been shown to be the target of virtually all lumbar spinal projection neurons of the anterolateral tracts in rodents [64]. To visualize spinoparabrachial neurons, the retrograde tracer CTB was stereotaxically injected into the lPBN on the side contralateral to the intraspinal viral injection (Figure 4d,g), and the number of hM3Dq-mCherry and CTB-labeled neurons were quantified across the dorsal horn (Figure 4e). Throughout the dorsal horn, a small percentage of hM3Dq-mCherry neurons (SDH: 16.7%, DDH: 1.9%, LSN: 6.7%) were CTB-backlabeled spinoparabrachial neurons, suggesting that *Tacr1*^*CreER*^ neurons are predominantly interneurons (Figure 4f). This observation is striking as it calls into question whether NK1R is an acceptable marker for spinal projection neurons [9,18,71]. While NK1R is expressed in spinal projection neurons, it is not exclusively expressed in spinal projection neurons.

Previous immunohistochemistry and *in situ* hybridization studies in wild-type mice estimate that between 65-90% of all mouse spinoparabrachial neurons express NK1R or *Tacr1*, respectively [3,18,25]. We report that a smaller percentage of CTB-backlabeled spinoparabrachial neurons (SDH: 32.1%, DDH: 12.9%, LSN: 16.7%) coexpressed hM3Dq-mCherry (Figure 4f). However, this observation is consistent with the efficiency of our genetic (*Tacr1*^*CreER*^ allele) and viral tools for targeting NK1R spinal neurons.

Lastly, to determine whether CNO administration preferentially activated one Tacr1::hM3Dq-mCherry neuron population over another, we quantified the percentage of hM3Dq-mCherry and hM3Dq-mCherry, CTB neurons that were immunoreactive for cFOS (Supp Figure 2). We found that cFOS expression was equivalent between hM3Dq-mCherry and hM3Dq-mCherry, CTB neurons, indicating that Tacr1::hM3Dq-mCherry interneurons and spinal projection neurons were equally activated by CNO.

### NK1R is expressed in GRPR interneurons within the superficial dorsal horn

Based on the observations that NK1R spinal neurons modulate itch and that the majority of *Tacr1*^*CreER*^ neurons manipulated in chemogenetic behavioral studies were interneurons, we wondered where NK1R interneurons fit within the current model of spinal itch circuitry. The current model of spinal itch transmission positions GRPR interneurons as a cellular gate of itch. Therefore, we asked whether NK1R is expressed in GRPR interneurons.

GRPR expression is restricted to neurons within the SDH [9,61,62], and thus coexpression analyses were similarly limited to the SDH in our studies. We first tested if NK1R and GRPR mRNA transcripts are expressed within the same neurons. Dual FISH studies on lumbar spinal cord slices from C57BL/6 mice demonstrated notable overlap between *Tacr1* and *Grpr*, with 48.1% ± 4.3 of *Tacr1* SDH cells coexpressing *Grpr*, and 38.8% ± 3.1 of *Grpr* SDH cells coexpressing *Tacr1* (Figure 5a,b).

**Figure 5.**
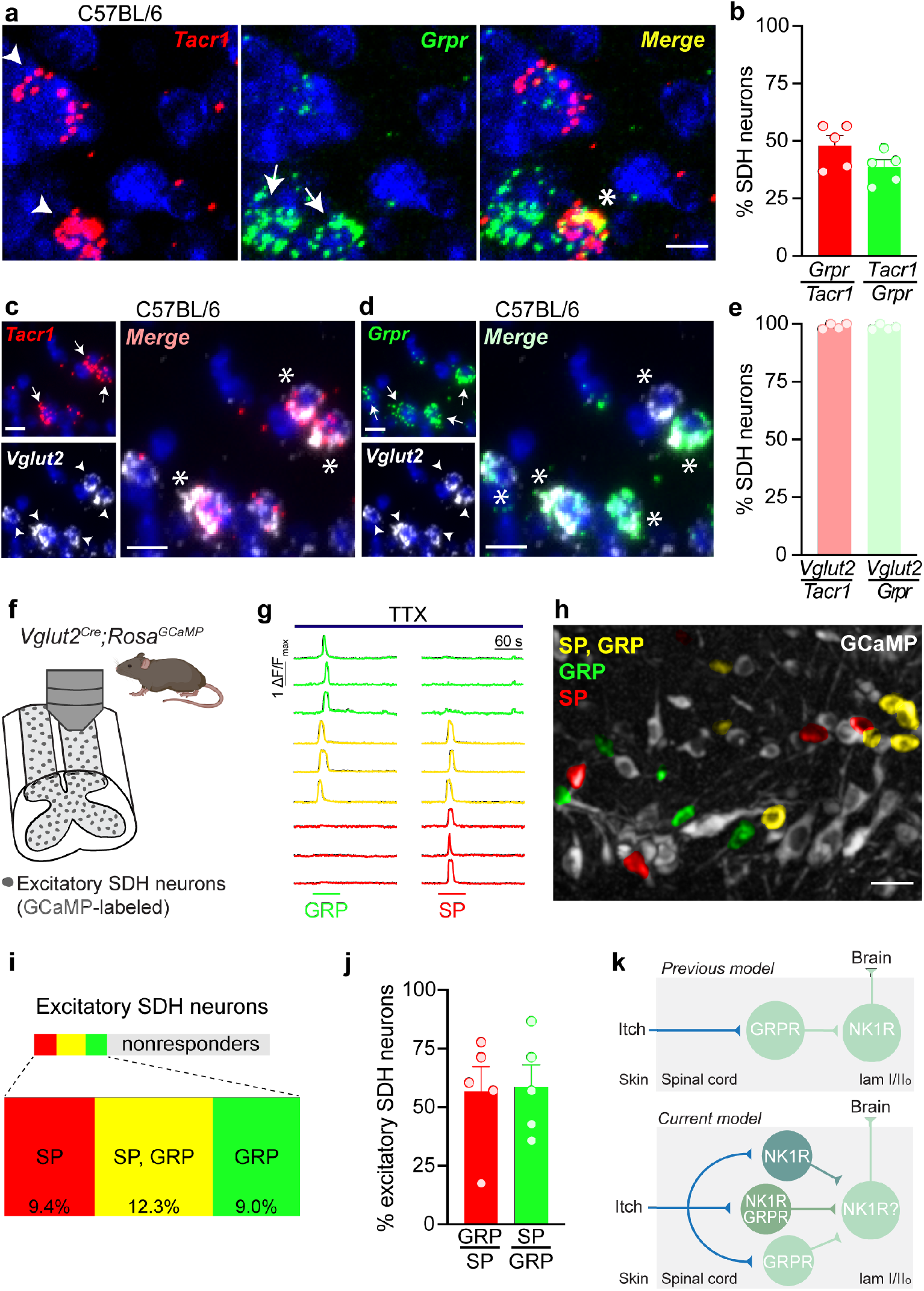
NK1R neurons in the superficial dorsal horn are a subset of GRPR interneurons. **a**, Representative images of dual FISH for *Tacr1* (red) and *Grpr* (green) performed on lumbar spinal cord sections from C57BL/6 mice. Scale bar, 10 μm. **b**, Percentage of SDH neurons that coexpress *Tacr1* and *Grpr* in C57BL/6 mice (n=5 mice). Representative images of dual FISH evaluating the expression of the excitatory neuronal marker *Vglut2* in **c**, *Tacr1*- and **d**, *Grpr*-expressing SDH neurons in C57BL/6 mice. Scale bars, 10 μm. **e**, Percentage of *Tacr1* and *Grpr* SDH neurons that coexpress *Vglut2* in C57BL/6 mice (n=4 mice). **f**, Schematic illustrating experimental set up for Ca^2+^ imaging of excitatory SDH neurons using *Vglut2*^*Cre*^;*Rosa*^*GCaMP6s*^ mice. **g**, Representative traces of Ca^2+^ transients in response to 1 μM SP and 300 μM GRP in the presence of 500 nM TTX. **h**, Representative psuedocolored fluorescent image showing *Vglut2*^*Cre*^;*Rosa*^*GCaMP6s*^ SDH neurons (grey) activated by SP (red), GRP (green), or both (yellow). Scale bars, 25 μm. **i**, Treemap of the percentage of excitatory SDH neurons that responded to SP (red), GP (green), both SP and GRP (yellow), or neither (grey) (n= 5 mice, 57-189 *Vglut2*^*Cre*^;*Rosa*^*GCaMP6s*^ neurons/mouse). **j**, Percentage of SDH neurons activated by both SP and GRP in *Vglut2*^*Cre*^;*Rosa*^*GCaMP6s*^ mice (n=5 mice). **k**, Schematic of the previous (top) and current (bottom) proposed model of the positions of NK1R and GRPR neurons within itch spinal circuitry. Solid lines do not necessarily represent direct synaptic connections. SDH, superficial dorsal horn; TTX, tetrodotoxin; SP, substance P; GRP, gastrin-releasing peptide. Data are shown as mean ± SEM, with open circles representing individual mice.

To address whether NK1R and GRPR are functionally coexpressed within the same neurons, we decided to utilize two-photon Ca^2+^ imaging within an *ex vivo* spinal cord preparation, as NK1R and GRPR are both Gq-coupled GPCRs that cause release of intracellular Ca^2+^ stores upon activation [13,35]. To determine the appropriate Cre driver for the genetically encoded Ca^2+^ indicator GCaMP6s, we performed dual FISH studies. These showed extensive colocalization of the glutamatergic neuronal marker *Vglut2* with *Tacr1* (99.0% ± 0.6 of *Tacr1* neurons) and *Grpr* (98.5% ± 0.5 of *Grpr* neurons) in SDH neurons, indicating that these are excitatory neuron populations (Figure 5c-e), in agreement with previous studies [20,25,71]. Therefore, we selectively targeted *Vglut2*-positive excitatory neurons within the SDH in Ca^2+^ imaging experiments. To assess functional expression of NK1R and GRPR in SDH neurons, Ca^2+^ imaging experiments were performed in the presence of TTX so that neurons that directly responded to SP or GRP, and thus express NK1R or GRPR, respectively, could be identified (Figure 5g,h).

In alignment with FISH analyses, we observed considerable overlap in responsivity to SP and GRP. Specifically, we found that out of all excitatory SDH neurons, 12.3% responded to both SP and GRP, while 9.4% responded to SP alone, and 9.0% responded to GRP alone (Figure 5i). Analysis of the excitatory SDH neurons that responded to either agonist revealed that 56.9% ± 10.5 of SP-responsive SDH neurons were also activated by GRP, and 58.8% ± 9.3 of GRP-responsive SDH neurons were also activated by SP (Figure 5j). Taken together, these results show that NK1R is functionally expressed within a subset of excitatory GRPR interneurons within the superficial dorsal horn. Importantly, this evidence demonstrates that NK1R is well-positioned to modulate spinal itch transmission.

## Discussion

The results of this study address two important questions regarding NK1R spinal dorsal horn neurons: 1) Is NK1R a selective marker for ascending spinal projection neurons? and 2) Where do NK1R neurons fit within the current model of spinal itch transmission?

### NK1R does not selectively mark spinal projection neurons

Although previous reports have acknowledged that NK1R is expressed in spinal projection neurons, as well as interneurons [21,63], NK1R is still frequently (and inappropriately) described as a selective marker for spinal projection neurons. Moreover, the presence or absence of NK1R immunoreactivity has been used as an inclusion criteria for determining whether or not a given dorsal horn neuron population includes spinal projection neurons [9,18,28,71].

Here, we demonstrated that *Tacr1* is endogenously expressed throughout the dorsal horn: in superficial and deeper dorsal horn neurons, as well as the lateral spinal nucleus. This spatial distribution was recapitulated in our *Tacr1*^*CreER*^ mouse line, and retrograde labeling from the parabrachial nucleus in these mice shows that projection neurons only represent a small fraction (<10%) of all *Tacr1* spinal neurons, highlighting that the majority of NK1R neurons are actually interneurons. Thus, while NK1R marks spinal projection neurons, it does not *selectively* mark spinal projection neurons.

The idea that NK1R selectively marks spinal projection neurons originates from a variety of circumstantial findings. For instance, early neurotoxic ablation studies have shown loss of NK1R immunoreactivity was largely limited to lamina I of the dorsal horn – where spinal projection neurons are most concentrated – likely due to rapid degradation and poor spread of SP-saporin [40,66]. Furthermore, the finding that NK1R immunoreactivity is lower in interneurons than spinal projection neurons [21] has been oversimplified, leading to the idea that NK1R immunoreactivity identifies projection neurons. Lastly, the use of NK1R as a selective projection neuron marker also stemmed from a lack of alternative spinal projection neuron markers. To date, NK1R remains the marker that captures the greatest proportion (an estimated 65-90% of projection neurons in mouse) [3,18,25], though recent single cell RNA-sequencing studies hold promise for novel spinal projection neuron markers, such as *Lypd1* [25]. Overall, while these studies have been essential in identifying NK1R as a potential marker and therapeutic target, the assertion that NK1R is a specific spinal projection marker represents an oversimplification of this previous work.

### A new role for NK1R interneurons in spinal itch transmission

NK1R spinal neurons have previously been implicated in itch, but further characterization of NK1R spinal neurons and how they fit within the current model of spinal itch transmission has remained unexplored. Here, we provide evidence that pharmacological activation of spinal NK1R and chemogenetic activation of *Tacr1*^*CreER*^ spinal neurons potentiates acute itch behavior, whereas pharmacological inhibition of spinal NK1R suppresses acute itch behavior, complementing previous NK1R neurotoxic ablation studies [1,3,4,19].

A key finding of the present study is that NK1R interneurons likely contribute to spinal itch transmission. We showed that the majority of *Tacr1*^*CreER*^ superficial dorsal horn neurons are interneurons, and that chemogenetic activation of these neurons increases behavioral responses to the pruritogen chloroquine. Further, our Ca^2+^ imaging findings indicate a subpopulation of excitatory neurons within the superficial dorsal horn functionally expresses NK1R, and that at least half of these neurons coincide with GRPR interneurons, which are a critical hub for spinal itch transmission [37,38,45,61,62].

One surprising observation from our study is that different strategies for activating NK1R spinal neurons produced different effects on spontaneous itch behavior. While pharmacological activation of spinal NK1R elicited robust spontaneous itch-related behaviors, chemogenetic activation of *Tacr1*^*CreER*^ spinal neurons did not. One possible explanation for this observation is that NK1R and hM3Dq – though both Gq-coupled GPCRs – engage different intracellular signaling pathways. Another striking finding was that chemogenetic activation of *Tacr1*^*CreER*^ spinal neurons increased itch, but not pain, behaviors. This finding might be due to the efficiency of the *Tacr1*^*CreER*^ allele, which captures about half of all *Tacr1* neurons, or possibly reflect that the allele preferentially targets a subset *Tacr1*^*CreER*^ spinal neurons that integrate itch.

### NK1R is expressed in a subpopulation of GRPR interneurons

Whether NK1R and GRPR are coexpressed within superficial dorsal horn neurons has been a controversial question. Previous histology studies on GRPR-eYFP mice reported that NK1R and GRPR are nonoverlapping markers [9]. In contrast, *Tacr1* and *Grpr* were found to demarcate an excitatory neuron population (“*Glut12*”) in a recent unbiased classification of dorsal horn neuronal subtypes [25,70]. Here, we provide compelling evidence using FISH and Ca^2+^ imaging that the two receptors are coexpressed at the both the transcript and functional receptor level. Likewise, our behavioral findings similarly point to functional overlap of NK1R and GRPR neurons in itch signaling, as the behavioral responses to intrathecal injection of SP, the selective NK1R agonist GR 73, 632, and GRP closely resembled one another. One interpretation of this finding is that these behaviors reflect the activation of a common neural substrate for itch (e.g. NK1R/GRPR interneurons) by the different agonists.

Importantly, the finding that NK1R is expressed within a subset of GRPR interneurons similarly positions NK1R in the center of spinal itch circuitry. Our results therefore suggest that the previous model of spinal itch transmission, which proposed a linear relationship between GRPR interneurons and NK1R spinal projection neurons, requires updating. We propose that spinal integration of itch input by NK1R and GRPR neurons is far more complex, with several interneuron subpopulations – those expressing NK1R, GRPR, or both NK1R and GRPR – integrating itch input, which is then likely relayed to the brain by NK1R spinal projection neurons (Figure 5k).

Our data also indicate that within the SDH, NK1R and GRPR are not entirely overlapping populations; rather, approximately half of each population overlaps with the other. This finding suggests that GRPR interneurons are a heterogenous population, as highlighted by recent electrophysiological characterization of GRPR-EFYP neurons [9,45]. Whether NK1R/GRPR neurons convey itch through the same or different mechanisms as GRPR neurons lacking NK1R, is an important question for future studies.

### NK1R as a therapeutic target for itch – What have we learned?

Our study provides evidence that NK1R spinal neurons mediate behavioral responses to itch in rodents. Taken together with additional studies showing that NK1R blockade and ablation reduce acute and chronic itch in rodent models [1,3,4,19], NK1R/GRPR interneurons are a plausible target for NK1R antagonists that were efficacious in reducing itch severity scores in Phase II clinical trials [46,48,49,58–60,69]. However, current NK1R antagonists appear to have limited clinical efficacy. The results of Phase III clinical trials are beginning to unfold, reporting that NK1R antagonist treatment failed to significantly improve itch ratings relative to placebo (NCT03540160). Thus, it is becoming apparent that while NK1R certainly plays a part in itch, its the role is likely a modulatory one.

It is interesting to compare the development of NK1R antagonists for the treatment of chronic pain and chronic itch. Analogous NK1R ablation and inhibition studies in rodents suggested NK1R as a therapeutic target for pain; however, NK1R antagonists failed broadly in clinical trials and NK1R was quickly abandoned as a therapeutic target [14,26]. NK1R antagonists have undoubtedly shown greater clinical efficacy in the treatment of chronic itch than chronic pain, yet are likely to be pulled once again from development. Although patients are itching for relief, the trials and tribulations of the development of NK1R as a therapeutic target for itch and pain may represent a cautionary tale that thorough preclinical testing is necessary before moving onto clinical trials.

Nonetheless, the findings of the present study highlight a new role for NK1R interneurons in itch, which comprise a subpopulation of excitatory GRPR interneurons. More broadly, this works adds to the current understanding of neurons that transmit itch within the spinal cord, which may in turn be leveraged for novel treatments in the future.

## Acknowledgements

We thank all members of the Ross lab for their comments and suggestions, as well as Michael C. Chiang and Justin Chestang for their technical assistance. Special thanks to Dr. Kelly Smith for advice on figure design and Dr. David Baranger for statistical consultation. Supported in part by National Institutes of Health grants T32NS086749 (TDS), F32NS110155 (TDS), T32 NS073548 (CAW), and R01NS096705 (SER).

## Author Contributions

Conceived and designed experiments: TDS, SER. Performed the experiments & collected data: TDS, CAW, LGF. Analyzed the data: TDS. Wrote the paper: TDS. All authors edited the manuscript.

