## Supplementary Data for "The neurokinin-1 receptor is expressed with gastrin-releasing peptide receptor in spinal interneurons and modulates itch"

### Supplemental Data

| *Comparison* | Nociceptive Threshold | *p* | *t* | *df* |
|---|---|---|---|---|
| *Baseline* Tacr1::control vs Tacr1::hM3Dq | Heat | 0.357 | 1.32 | 32 |
| | Cold | 0.846 | 0.338 | 32 |
| | Mechanical | 0.619 | 0.885 | 32 |

**Supplementary Table 1**. Statistical analyses of baseline nociceptive withdrawal thresholds of the ipsilateral hindpaw in chemogenetic behavioral studies (related to Figure 3). Post hoc t-Tests, Holm-Sidak correction for multiple comparisons (n=8-10 mice/group).

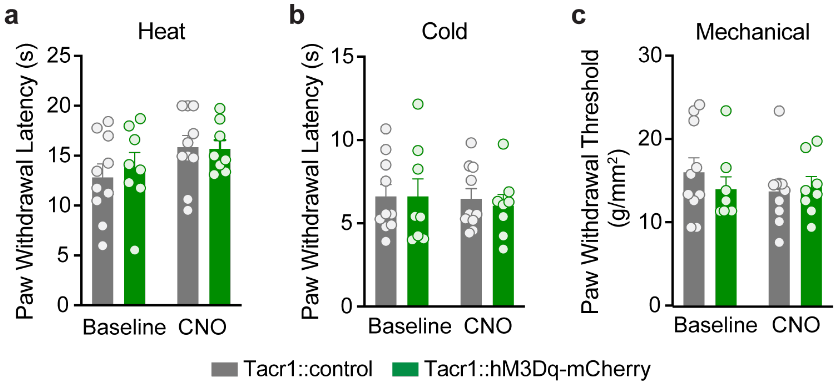


**Supplementary Figure 1.** Chemogenetic activation of *Tacr1*$^{CreER}$ spinal neurons does not affect nociceptive withdrawal thresholds of the hindpaw contralateral to intraspinal viral injection (related to Figure 3). Chemogenetic activation of *Tacr1*$^{CreER}$ neurons with CNO administration (5 mg/kg, i.p.) did not affect the contralateral hindpaw withdrawal thresholds of Tacr1::hM3Dq-mCherry mice to **a,** heat (2-way RM ANOVA, CNO x DREADD, $p$=0.699, $F$ (1,16)=0.155, n=8-10 mice/group), **b,** cold (2-way RM ANOVA, CNO x DREADD, $p$=0.753, $F$ (1,16)=0.102, n=8-10 mice/group), or **c,** mechanical (2-way RM ANOVA, CNO x DREADD, $p$=0.0.338, $F$ (1,16)=0.789, n=8-10 mice/group) stimuli relative to Tacr1::controls. Data are shown as mean ± SEM, with open circles representing individual mice.

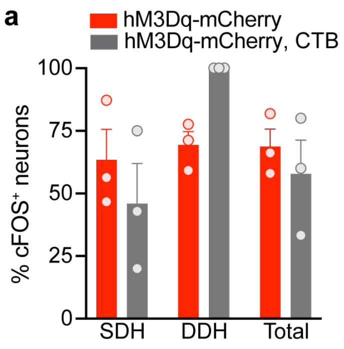


**Supplementary Figure 2.** CNO administration activates *Tacr1*$^{CreER}$ interneurons and spinoparabrachial neurons equally (related to Figure 4). Data are the same as those reported in Figure 3k-l, but are reanalyzed here to evaluate cFOS immunoreactivity in interneurons versus spinoparabrachial neurons of Tacr1::hM3Dq-mCherry mice. **a,** Quantification of the percentage of cFOS$^{+}$ hM3Dq-mCherry and hM3Dq-mCherry, CTB neurons across the dorsal horn shows equal activation following CNO administration (Student's t-Test, $p$=0.513, t=7.18, df=4, n=3 mice). cFOS expression was detected within a single LSN in one mouse (Figure 3l), and thus LSN neurons are not included here. Data are shown as mean ± SEM, with open circles representing individual mice.
